# A new framework for analysis of three-dimensional shape and architecture of human skeletal muscles

**DOI:** 10.1101/2021.09.08.459536

**Authors:** Bart Bolsterlee

## Abstract

A new framework is presented for comprehensive analysis of the three-dimensional shape and architecture of human skeletal muscles from magnetic resonance and diffusion tensor imaging data. The framework comprises three key features: (1) identification of points on the surface of and inside a muscle that have a correspondence to points on and inside another muscle, (2) reconstruction of average muscle shape and average muscle fibre orientations, and (3) utilization of data on between-muscle variation to visualize and make statistical inferences about changes or differences in muscle shape and architecture. The general use of the framework is demonstrated by its application to three datasets. Analysis of data obtained before and after eight weeks of strength training revealed there was little regional variation in hypertrophy of the vastus medialis and vastus lateralis, and no systematic change in pennation angle. Analysis of passive muscle lengthening revealed heterogeneous changes in shape of the medial gastrocnemius, and confirmed the ability of the methods to detect subtle changes in muscle fibre orientation. Analysis of the medial gastrocnemius of children with unilateral cerebral palsy showed that muscles in the more-affected limb were shorter, thinner and less wide than muscles in the less-affected limb, and had slightly more pennate muscle fibres in the central and proximal part of the muscle. Amongst other applications, the framework can be used to explore the mechanics of muscle contraction, investigate adaptations of muscle architecture, build anatomically realistic computational models of skeletal muscles, and compare muscle shape and architecture between species.

## Introduction

A remarkable property of muscles is that they can atrophy and hypertrophy rapidly in response to mechanical loading, such as the unloading that occurs with bed rest (LeBlanc et al., 1992; Marusic et al., 2021) or the increased loading that occurs with strength training (Jones and Rutherford, 1987; Jorgenson et al., 2020). Changes in muscle volume are the result of an imbalance between contractile protein turnover and degradation (Sartori et al., 2021), and must reflect a change in the size and arrangement (architecture) of muscle fibres. Muscle fibre size and architecture are thought to be the primary determinants of the functional capacity of muscles (Otten, 1988; Powell et al., 1984). Consequently, adaptations of muscle architecture have been studied extensively (e.g. Aagaard et al., 2001; Blazevich, 2019; Erskine et al., 2010; Franchi et al., 2014; Gillett et al., 2016; Kawakami et al., 1995).

Muscle hypertrophy and atrophy are typically quantified by changes in muscle volume, cross-sectional area or thickness (Franchi et al., 2018). These scalar metrics may be obtained from two-dimensional (2D) ultrasound imaging or 3D magnetic resonance imaging (MRI). Muscle architecture (fascicle lengths and pennation angles) is usually measured in 2D from ultrasound images (Blazevich et al., 2007; Erskine et al., 2010). A limitation of these scalar metrics is that they do not characterise the complex 3D muscle shape and architecture of human skeletal muscles (e.g. Agur et al., 2003; Bolsterlee et al., 2018; Lee et al., 2015). As a result, limited information is available on how skeletal muscles adapt in 3D, and what the functional consequences of these adaptations are.

Diffusion tensor imaging (DTI) is an MRI protocol from which 3D local muscle fibre orientations can be derived. DTI reconstructions of skeletal muscle architecture are based on the principle that water molecules in muscle tissue diffuse over longer distances along the muscle fibre than in its transverse plane (Damon et al., 2002; Schenk et al., 2013). Previously we have shown that DTI can provide reliable measurements of the average fascicle length of human muscles (Bolsterlee et al., 2019), but measurements of average fascicle length do not exploit the full potential of DTI to quantify muscle architecture across a whole muscle. One obstacle to accurate local measurements of muscle architecture is the relatively low signal-to-noise ratio of DTI scans, sometimes leading to sparse reconstructions with regions where no or inaccurate measurements of fibre orientations are obtained. More accurate and complete muscle architecture reconstructions can potentially be obtained by averaging across scans from different individuals. However, building group-averaged models of fibre architectures requires averaging of diffusion tensors across participants, which is not as straightforward as averaging scalar metrics. Arsigny et al. (2006) laid out the foundations for mathematical operations like averaging and interpolation of diffusion tensors, and that work was subsequently used by Peyrat et al. (2007) to build population-averaged atlases of 3D fibre architectures of canine and human hearts. The methods used in those studies can form the basis for comprehensive analyses of architectural adaptation of skeletal muscles.

The aim of this manuscript is to describe new methods that can be used to comprehensively describe, visualise, quantify and statistically analyse 3D changes in skeletal muscle shape and architecture. To this end, a new framework for building group-averaged models of skeletal muscle shape and architecture is outlined. Data from a strength training study (Bolsterlee et al., 2021) are used to describe the methods in detail, and to illustrate the use of these methods in muscle hypertrophy studies. To further test and demonstrate the general use of these methods, they are also applied to analyse shape and architecture during passive muscle lengthening (Bolsterlee et al., 2017) and to compare muscles between the more- and less-affected side of children with unilateral cerebral palsy (D’Souza et al., 2019).

## Methods

### Dataset 1: Strength training

The methods are described by applying them to the analysis of muscle hypertrophy, using data from a study which investigated the effect of strength training on intramuscular fat in human thigh muscles. A report of that study, which includes participant characteristics and the training protocol, has been published elsewhere (Bolsterlee et al., 2021). Briefly, 11 healthy young adults trained their knee extensor muscles three times per week for eight weeks with leg extension and leg press exercises. MRI and DTI scans were obtained before and after the training period (see Appendix 1 for imaging protocol). Compared to before training, isometric knee extensor strength after training was 12 ± 14% greater (mean ± S.D. across participants), and the vastus medialis and vastus lateralis were 14 ± 9% and 15 ± 10% larger in volume, respectively.

### Muscle shape analysis

Using ITK-SNAP (Yushkevich et al., 2006), the vastus medialis and vastus lateralis muscles were manually segmented from the mDixon scans. From the segmentations 3D triangulated surface models with edge lengths of ∼5 mm (Fig. 1A) were created using the Matlab iso2mesh toolbox (The Mathworks Inc.; Natick, Massachusetts, USA; version 2019b) (Fang and Boas, 2009). The mDixon scan did not cover the most proximal part of the muscles, so surface models were clipped in the transverse plane at 70% of the distance from the lateral epicondyle to greater trochanter.

**Figure 1.**
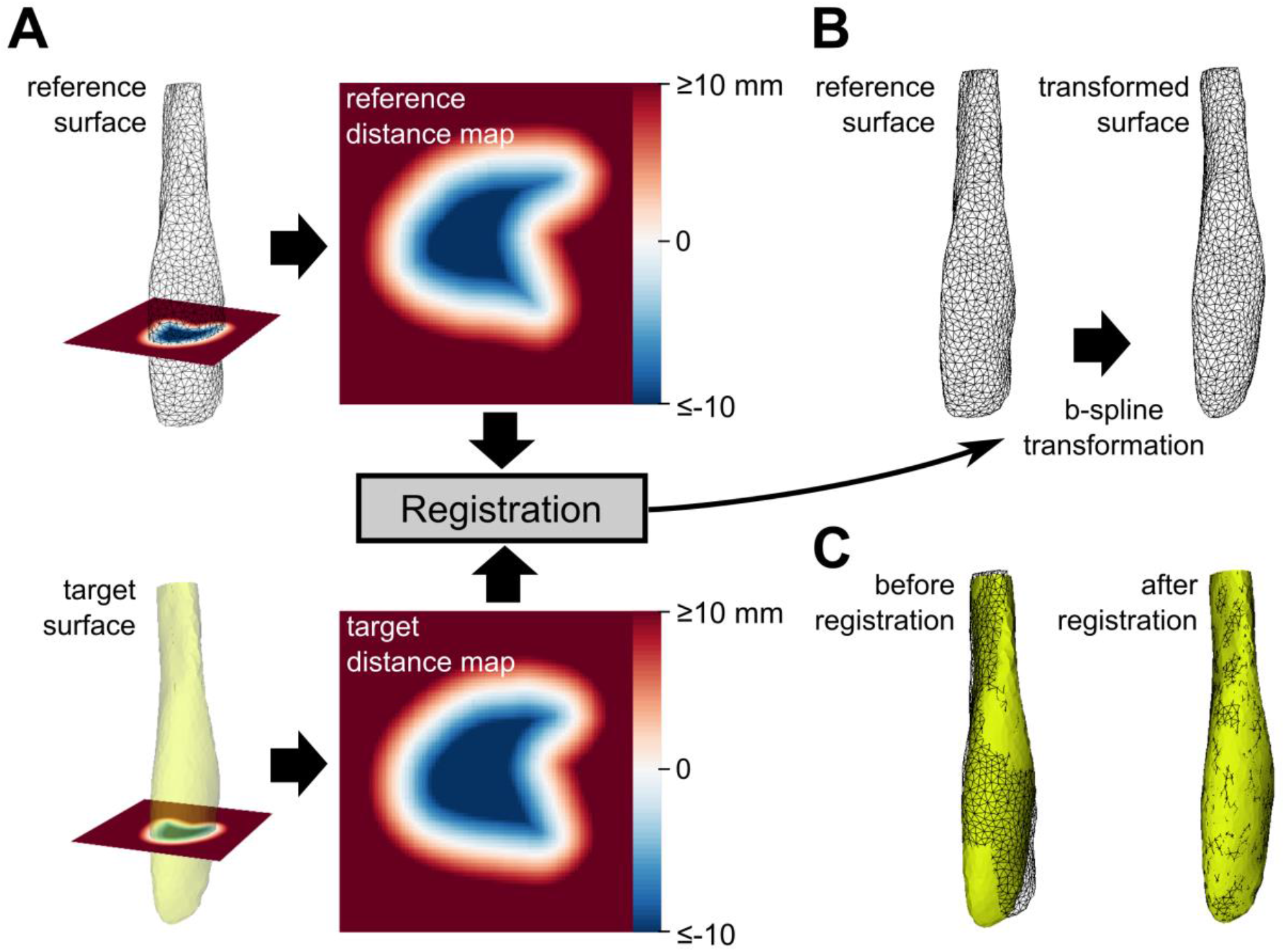
A) Illustration of non-rigid registration of distance maps to establish point-to-point correspondence between surfaces. Distance maps of a reference surface (top; black wireframe) and a target surface (bottom; yellow) were registered using a b-spline transformation. B) The b-spline transformation was then applied to the vertices of the reference surface. C) Overlay of the target surface (yellow) and the original (left) and transformed (right) reference surface, showing that the transformed surface matched the target surface well.

Building group-averaged 3D muscle shapes required finding corresponding vertices on the surface of all muscles (Heimann and Meinzer, 2009). First, muscle surfaces from 10 of the 11 participants were rigidly aligned to the surface of the muscle from the remaining participant (the “reference” muscle) using the iterative closest point algorithm built into ShapeWorks (Cates et al., 2017). For all rigidly aligned muscles, distance maps (resolution: 1×1×1 mm) were then created. A distance map is an image in which the value of a voxel is the smallest distance to the surface. Distance maps of all muscles were registered to the distance map of the reference muscle with a non-rigid b-spline transformation (grid size 20×20×20 mm; Elastix v4.7 (Klein et al., 2010)). The resulting transformation was then applied to all vertices of the reference muscle (1,333 vertices for the vastus medialis and 1,628 for the vastus lateralis) to obtain the corresponding vertices of all other muscles. All muscles were registered to a reference twice: in the first iteration, the muscle with the volume closest to the group-averaged volume was used as a reference; in the second iteration the mean shape from the first iteration was used as a reference to reduce bias towards the initially selected reference surface.

Registration of distance maps yields the transformation that optimally aligns locations that are equidistant from the surface (including points *on* the surface), which here we assume to be a reasonable solution to the ill-defined problem of point-to-point correspondence in muscle surfaces and volumes. The quality of registration of the muscle surface was assessed by interpolating the absolute distance map of the target surface at the vertices of the transformed reference surface. If the transformed surface perfectly represents the target surface, this distance, or the residual error, will be zero. The residual error after registration ranged from 0.4 to 0.8 mm for all 22 muscles, indicating that the transformed reference surface closely resembled the target surface.

The set of corresponding vertices on the surface of all muscles was used to calculate the average location of each vertex, and therefore the average shape across participants of each muscle both before and after training. The effect of training on muscle shape was defined as the nearest signed distance from the vertices of the average muscle surface after training to the average surface before training. Negative and positive values indicate that a vertex of the muscle after training is located inside (local contraction) and outside (local expansion) the surface before training, respectively (Fig. 1A). Note that these distances indicate the point-to-surface distance (smallest distance of a point to *any* point on the surface) rather than the point-to-point distance (distance between corresponding points). A test of the methods demonstrated that point-to-surface distances were estimated more accurately than point-to-point distances (Appendix 2). The vastus medialis and vastus lateralis surface models were colour-coded to visualise changes in shape with strength training (Fig. 3).

Bootstrapping was used to determine the sampling distribution of the distance between surfaces before and after training. In each of 1,000 bootstrap replicates of the original data sets, we sampled with replacement 11 pairs of muscles (before and after training) from 11 randomly selected participants, and then recalculated the mean shape before and after training. For all 1,000 iterations, the nearest distance from vertices of the mean surface after training to the mean surface before training was determined. The distribution of distances was corrected for sampling bias by adjusting the sampling distribution by a constant, so that for each vertex the mean of the distribution equalled the distance of the original dataset. Mean local expansion / contraction was deemed to be statistically significant at the 0.05 level if the 2.5^th^ / 97.5^th^ percentile of the sampling distribution was larger / smaller than 0, respectively, and not significant otherwise.

### Muscle architecture analysis

The DTI data were used to determine fibre orientations in all muscles, by assuming that the primary eigenvector of a diffusion tensor is aligned with local muscle fibre orientations (Damon et al., 2002; Froeling et al., 2012). Diffusion-weighted images were corrected for eddy-current induced distortions (Andersson and Sotiropoulos, 2016) and filtered with a nonlocal principal component analysis filter (Manjon et al., 2013). For each voxel the diffusion tensor was reconstructed, from which the eigenvectors and eigenvalues were extracted (DSI Studio; Yeh et al., 2013).

To determine the effect of training on group-averaged 3D fibre orientations, it is necessary to identify locations (nodes) inside muscles that correspond between muscles. Corresponding nodes inside the muscle volume were found by first defining a regular grid of nodes with a spacing of 5 mm inside the group-averaged muscle surface, and then applying the previously determined b-spline transformation to find those nodes in all other muscles (Fig. 2A).

**Figure 2.**
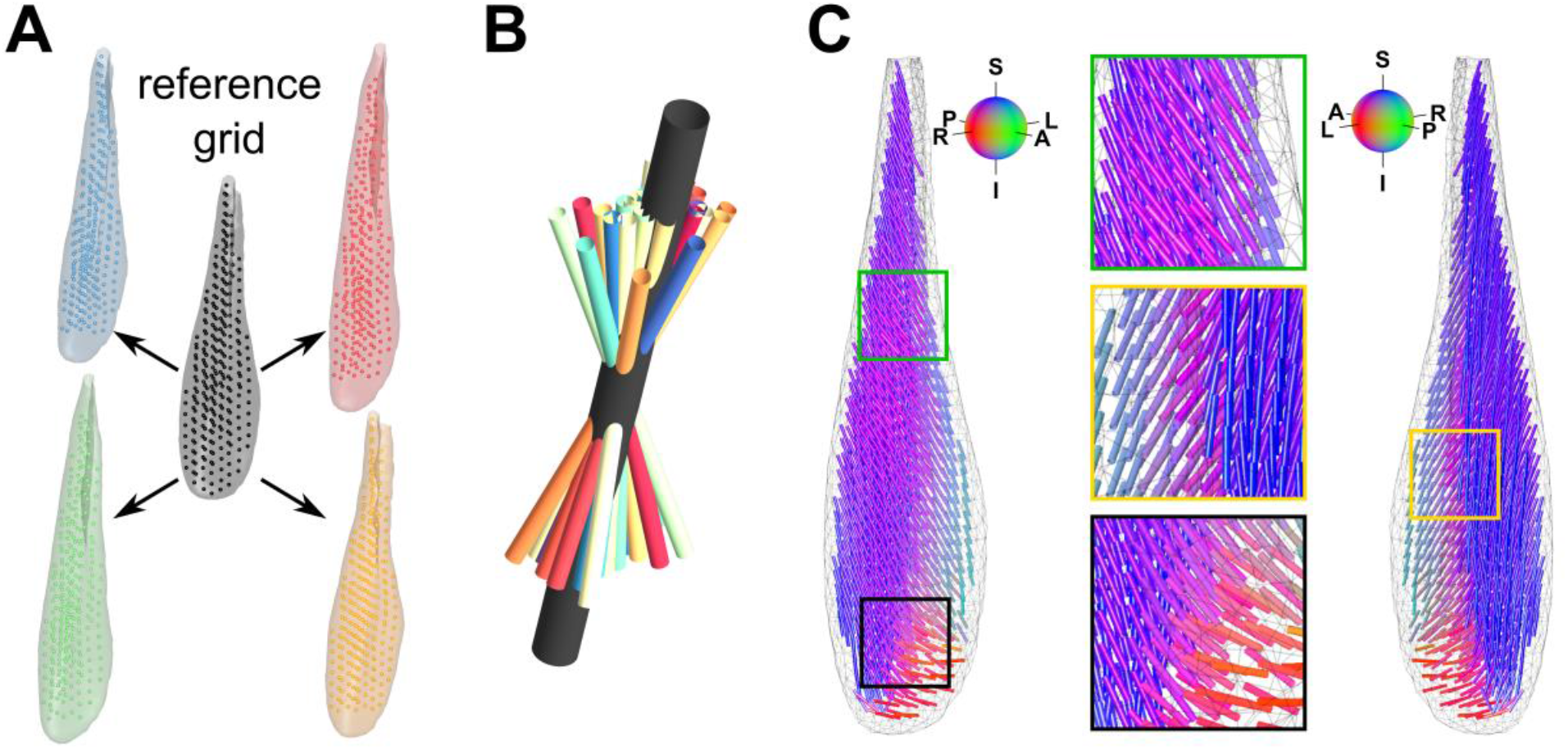
**A)** Illustration of how corresponding points inside muscle volumes were found. A regular grid of points inside the reference muscle (black) was transformed to each other muscle (coloured surfaces) using a b-spline transformation (see main text for details). B) Example of fibre orientations from individual muscles (coloured cylinders) and the group-averaged fibre orientation (black cylinder) obtained at one grid point, using averaging of diffusion tensors in the log-Euclidean framework. C) Two views on the reconstruction of the group-averaged fibre orientations of the vastus medialis. Fibres are coloured according to their direction (see colour spheres for interpretation). A=anterior, P=posterior, L=left, R=right, I=inferior, S=superior.

Next, fibre orientations were averaged across corresponding nodes. Sampling fibre orientations at locations other than voxel centroids and averaging fibre orientations across participants requires interpolation and averaging of diffusion tensors, which cannot simply be done directly on the components of the tensors (Arsigny et al., 2006; Yang et al., 2012). Instead, the log-Euclidean framework was used (Arsigny et al., 2005, 2006). In this framework, the 3×3 diffusion tensor for each voxel, *S*, is transformed to its log-Euclidean counterpart log(*S*), by:

1. diagonalising the diffusion tensor *S* into a rotation matrix *R* (a matrix of the eigenvectors of *S*) and a diagonal matrix 𝛬 (a diagonal matrix of the eigenvalues of *S*), so that *S* = *R*^*T*^ ∙ 𝛬 ∙ *R*;
2. calculating *log(*𝛬) by taking the natural logarithm of the diagonal components of 𝛬 (the eigenvalues of *S*);
3. recomposing to obtain *log*(*S*) = *R*^*T*^ ∙ *log(*𝛬) ∙ *R*.

Diffusion tensors are symmetric matrices with 6 degrees of freedom, so their 9 components can be represented in minimal form by a 6-element vector:

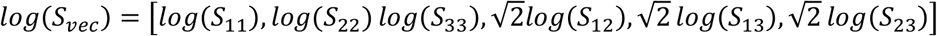

where *log*(*S*_*ij*_) is the (*i, j*)-th element of log(*S*).

Mathematical operations like interpolation and averaging can be conducted directly on the elements of log(*S*_*vec*_) (Arsigny et al., 2006). For all muscles, values for *log*(*S*_*vec*_) were interpolated at each node of the (transformed) grid, and then rotated with the previously determined rigid transformation to align all fibre orientations into a common coordinate system where the x-, y- and z-axes are the left-right, anterior-posterior and inferior-superior axes, respectively. The *log*(*S*_*vec*_) fields were then averaged across groups of muscles obtained before and after training. The averaged log(*S*_*vec*_) values were then transformed back from log-Euclidean space to the (group-averaged) diffusion tensor in Euclidean space, using the inverse of the operations described above. To determine the average fibre orientation at a node, the primary eigenvector was extracted using eigenvalue decomposition of the group-averaged diffusion tensor. Figure 2B shows an example of individual and group-averaged fibre orientations and Figure 2C shows a group-averaged muscle architecture reconstruction.

Due to noise in the DTI scans, realistic diffusion tensors could not be reconstructed at some nodes in each individual muscle. Nodes were excluded from calculation of the group-averaged values if eigenvalues of the diffusion tensor were negative, the mean diffusivity was more than two standard deviations away from the group-averaged mean diffusivity, or the fractional anisotropy was smaller than 0.1 (i.e. if diffusion was nearly isotropic).

To determine the effect of training on fibre orientations, the absolute angle in 3D between fibre orientations, averaged across participants, before training and after training were calculated. A second part of the analysis of the effect of training on fibre orientations focused on the pennation angle, which we defined as the acute angle between the superior-inferior axis (z-axis) and the (local) fibre orientation vector. (In other words, the pennation angle was defined as the inverse cosine of the absolute z-component of the primary eigenvector.) Pennation angles were determined for all individual muscles and nodes both before and after training. For all nodes, the average pennation angle before and after training, and the average change in pennation angle with training (subtracting values after training from values before training) were determined. As in the analysis of changes in muscle shape described above, bootstrapping was used to determine the sampling distribution and statistical significance of the change in pennation angle with training.

### Dataset 2: Passive muscle lengthening

To demonstrate the sensitivity of the methods to detect relatively small changes in 3D shape and architecture, the methods were applied to data from a previously published study on passive lengthening of the medial gastrocnemius muscle (Bolsterlee et al., 2017). In this study, anatomical MRI and DTI scans were obtained from the left calves of eight participants at three ankle joint angles at which the medial gastrocnemius was at its shortest and longest *in vivo* length, and at one intermediate length. The change in group-averaged 3D shape and pennation angle with passive lengthening was quantified, visualised and statistically analysed.

### Dataset 3: Cerebral palsy-related muscle adaptation

To demonstrate the use in studies of muscle architectural adaptations with disease, the methods were applied to data from a previously published study on the architecture of the medial gastrocnemius muscles of children with unilateral cerebral palsy (D’Souza et al., 2019). Here, we only use data from children with cerebral palsy (*n*=19, age range 5.4 to 15.8 years) and not from the typically developing children included as a control group in the original study. Muscle shape and fibre orientations from the left leg were mirrored in the sagittal plane to allow grouping with muscles from the right leg. Because of the small size of the children’s muscles, fibre orientations were sampled at nodes spaced 3 mm apart, instead of 5 mm for the adult muscles in the other datasets. The difference in 3D shape and architecture of the medial gastrocnemius between the more-affected and less-affected leg of children with unilateral cerebral palsy was quantified, visualised and statistically analysed.

### Code availability

Matlab algorithms to conduct all analyses, including example scripts and example data, are made available on GitHub (https://github.com/bartbols/MUSHAR-toolbox).

## Results

Group-averaged muscle shapes (Fig. 3) and fibre architectures (Fig. 4) were successfully reconstructed for all muscles in all datasets. Group-averaged fibre architecture reconstructions were substantially more complete than reconstructions from individual muscles. In the three datasets analysed, unrealistic diffusion tensors were found for, on average, 10 to 18% of nodes, and up to 65% in individual muscles. In contrast, all group-averaged reconstructions were complete or nearly complete (Table 1). The group-averaged reconstructions had smooth fibre orientations which clearly displayed known architectural features of the muscles, such as the oblique orientation of muscle fibres of the vastus medialis at its distal end (Fig. 4B) and the rather uniform orientation of muscle fibres in the vastus lateralis (Fig. 4D).

**Figure 3.**
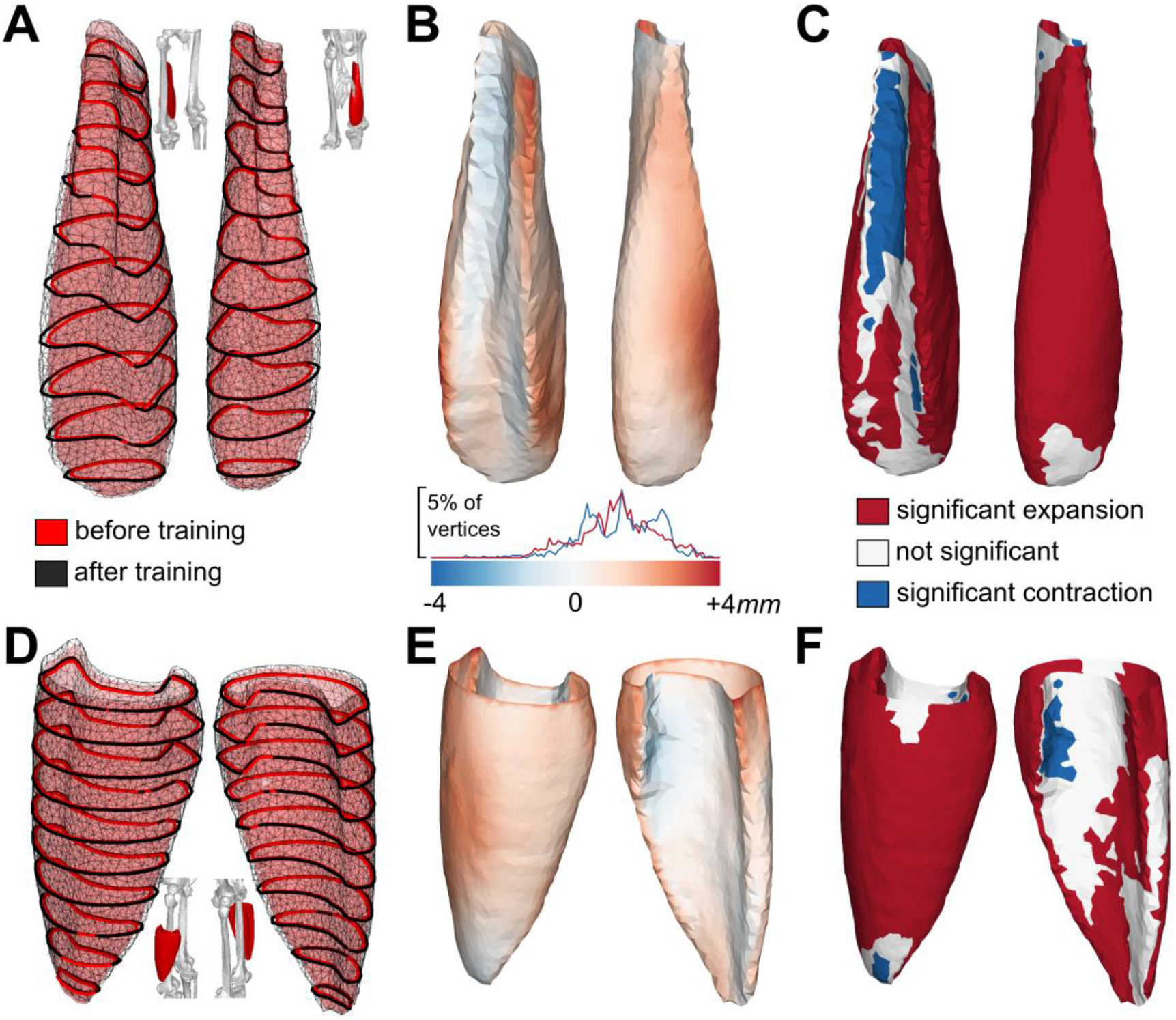
Effect of training on muscle shape of the vastus medialis (A-C) and vastus lateralis (D-F). Each panel shows the muscle viewed from two aspects. The left panels (A,D) show the group-averaged surface of muscles before (red transparent) and after training (black wireframe). Red and black lines are intersections with the transverse plane at 10% intervals along the muscle’s length. In B and E, the muscle surface is coloured according to the distance between the muscle surfaces before and after training (red/blue = local expansion/local contraction). The distribution of distances is plotted on top of the colour bar (red line for vastus medialis and blue line for vastus lateralis). In C and F, the muscle surfaces are coloured according to the statistical significance of the effect of training on muscle shape (red = significant expansion, white = no significant change, blue = significant contraction).

**Figure 4.**
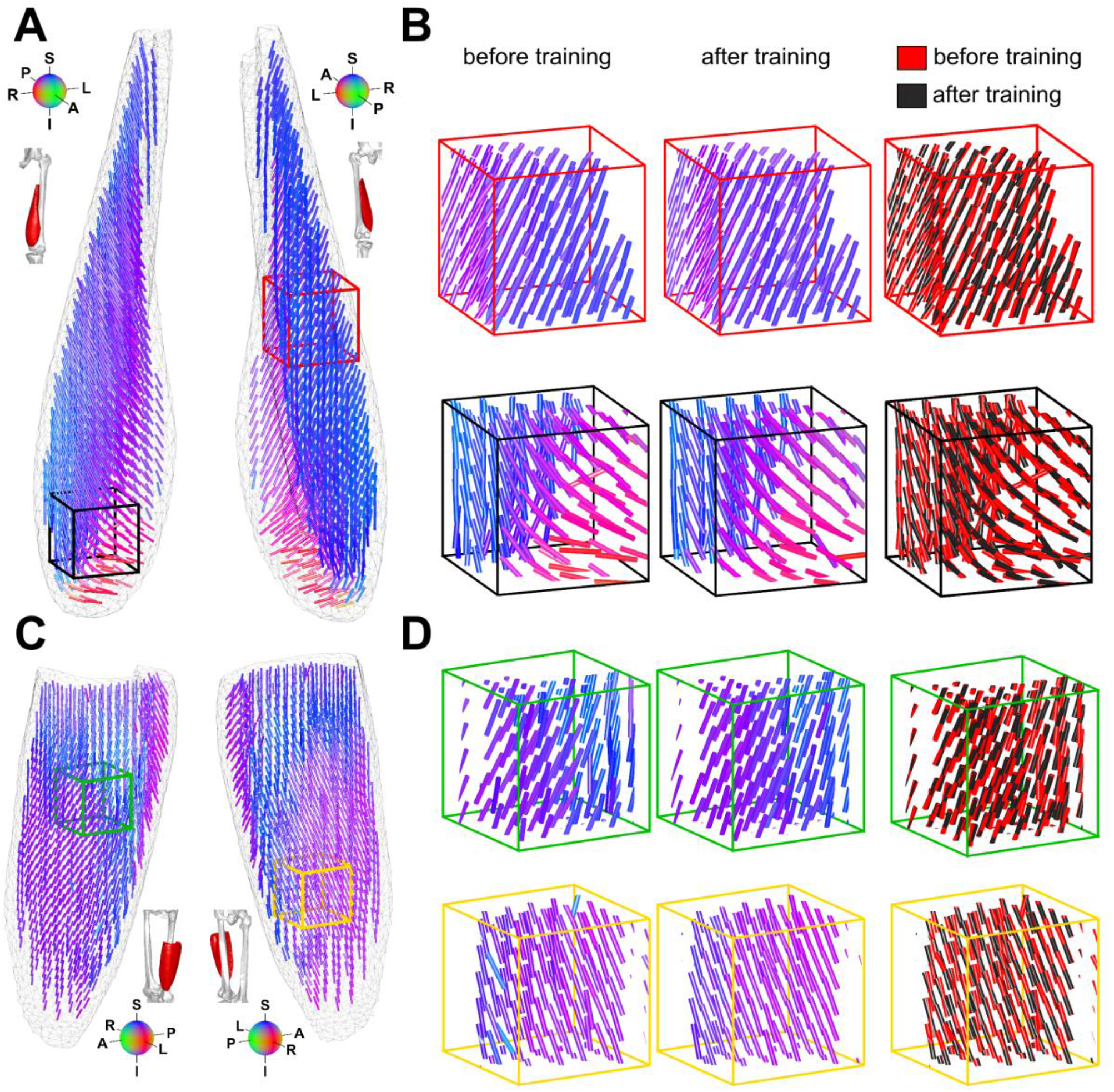
Effect of training on fibre orientations of the vastus medialis (A,B) and vastus lateralis (C,D). The left panels (A,C) show two views on the group-averaged fibre orientations before training. Panels B and D show zoomed in versions of 3 × 3 × 3 cm cubes inside the muscles (see A and C for locations) with, from left to right, fibre orientations before training, after training and an overlay of before (red) and after training (black). Fibres are coloured according to their direction (see colour spheres for interpretation). A=anterior, P=posterior, L=left, R=right, I=inferior, S=superior.

**Table 1.**
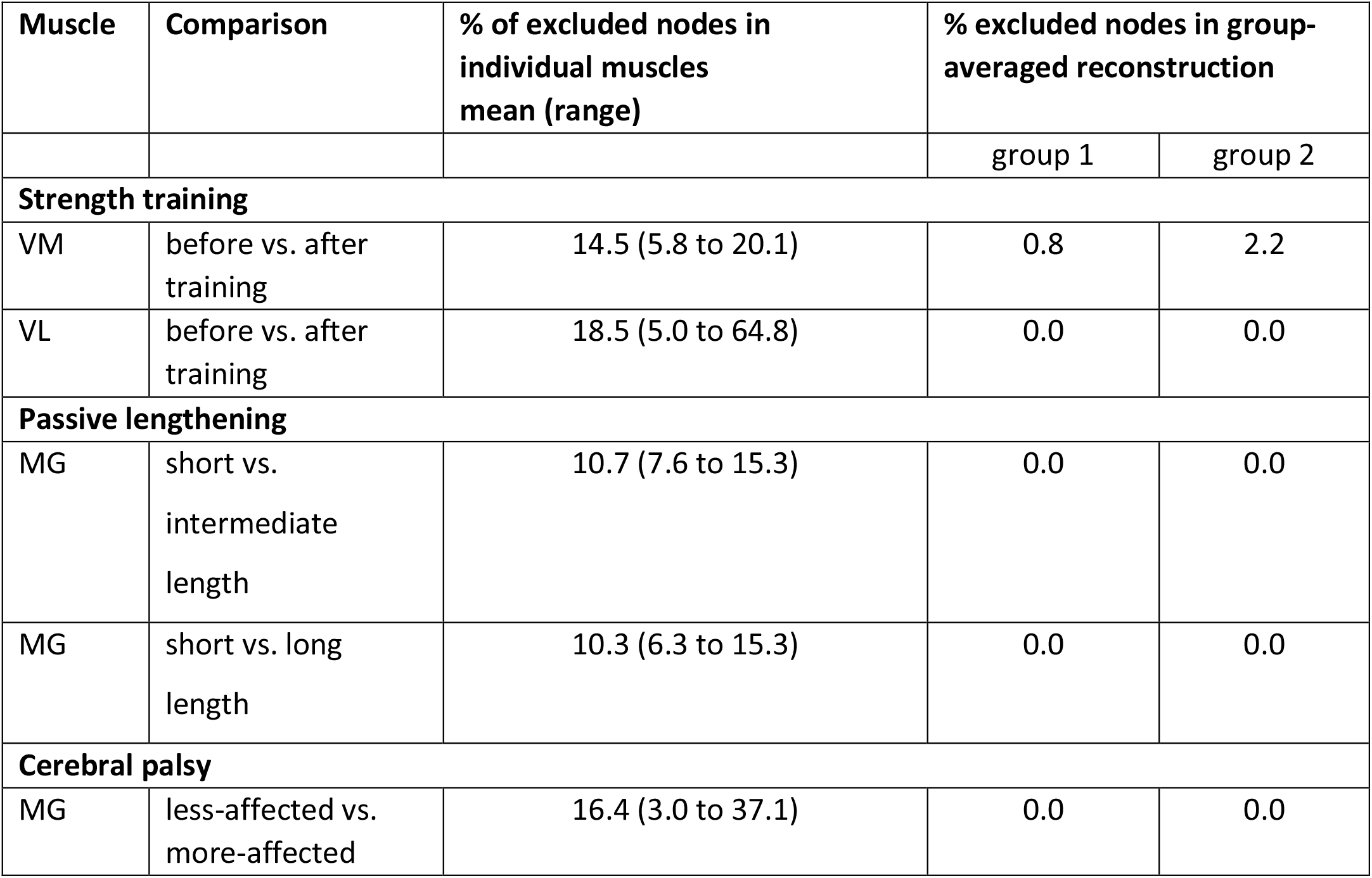
Overview of excluded nodes in individual muscle and group-averaged fibre architecture reconstructions. VM = vastus medialis, VL = vastus lateralis, MG = medial gastrocnemius

### Effect of training

Training resulted in an expansion of the muscle surface by, on average across vertices, 1.2 mm in both the vastus lateralis and vastus medialis (Table 2, Fig. 3). The increase in size was evident and statistically significant (Fig. 3D and F) everywhere along the muscles’ superior-inferior axis, mostly on the superficial surface (Fig. 3B and E). On the deep surface, the change in shape was small and often not statistically significant. The local expansion was statistically significant for 68-70% of vertices in the vastus medialis and vastus lateralis (Table 2).

**Table 2.** Overview of changes in muscle shape with strength training, passive lengthening and cerebral palsy. In the strength training dataset, results are also shown for the comparison between four muscles scanned twice on subsequent days before training (repeatability). *n* = number of muscles per group, VL = vastus lateralis, VM = vastus medialis, MG = medial gastrocnemius, N.S. = not significant, S.D. = standard deviation.

| Muscle | *n* | # of vertices | Comparison | distance (mm)<sup>§</sup><br>mean (S.D.) | Significance (% of vertices) |  |  |
| --- | --- | --- | --- | --- | --- | --- | --- |
|  |  |  |  |  | contraction | N.S. | expansion |
| **Strength training** |  |  |  |  |  |  |  |
| VM | 11 | 1,333 | before vs.<br>after training | 1.2 (1.0) | 6.7 | 25.6 | 67.7 |
| VL | 11 | 1,628 | before vs.<br>after training | 1.2 (1.1) | 2.8 | 27.4 | 69.7 |
| **Passive lengthening** |  |  |  |  |  |  |  |
| MG | 8 | 1,220 | short vs.<br>intermediate<br>length | 0.1 (0.7) | 29.1 | 41.1 | 29.8 |
| MG | 8 | 1,220 | short vs. long<br>length | 0.1 (1.4) | 32.9 | 38.5 | 28.6 |
| **Unilateral cerebral palsy** |  |  |  |  |  |  |  |
| MG | 19 | 1,187 | less-affected<br>vs. more-affected | -1.9 (1.5) | 77.6 | 13.0 | 9.5 |

*§ 'distance' is the nearest signed distance of a vertex on the mean surface of group 1 to the mean surface of group 2 (positive is local expansion, negative is local contraction).*

The change in 3D angle between group-averaged fibre orientations before and after training was 7.2 ± 7.1° in the vastus medialis and 11.9 ± 9.1° in the vastus lateralis (values are mean ± standard deviation across nodes). Based on visual (Fig. 4) and statistical analyses (Fig. 5, Table 3), we found no systematic effect of training on pennation angle. In both muscles, the change in pennation angle was not statistically significant in 87% of nodes (Table 3). The 13% of nodes where the change reached statistical significance did not display a consistent pattern (Fig. 5B and D).

**Table 3.** Overview of changes in pennation angle with strength training, passive muscle lengthening and cerebral palsy. In the strength training dataset, results are also shown for the comparison between four muscles scanned twice on subsequent days before training (repeatability). *n* = number of muscles, VL = vastus lateralis, VM = vastus medialis, MG = medial gastrocnemius, N.S. = not significant, S.D. = standard deviation.

| Muscle | *n* | # of nodes | Comparison | pennation (°)<br>mean (S.D.) |  |  | Significance<br>(% of nodes) |  |  |
| --- | --- | --- | --- | --- | --- | --- | --- | --- | --- |
|  |  |  |  | group 1 | group 2 | Δ | decrease | N.S. | increase |
| Strength training |  |  |  |  |  |  |  |  |  |
| VM | 11 | 1,177 | before vs.<br>after training | 26.3<br>(10.8) | 25.7<br>(10.6) | -0.2<br>(5.1) | 6.5 | 87.0 | 6.5 |
| VL | 11 | 1,082 | before vs.<br>after training | 29.0<br>(5.9) | 28.0<br>(6.1) | -2.7<br>(6.1) | 10.9 | 86.7 | 2.4 |
| Passive lengthening |  |  |  |  |  |  |  |  |  |
| MG | 8 | 1,438 | short vs.<br>intermediate<br>length | 24.1<br>(7.0) | 20.4<br>(7.7) | -3.6<br>(4.8) | 36.5 | 61.7 | 1.8 |
| MG | 8 | 1,438 | short vs.<br>long length | 24.1<br>(7.0) | 17.5<br>(8.5) | -6.6<br>(6.5) | 53.0 | 44.6 | 2.4 |
| Unilateral cerebral palsy |  |  |  |  |  |  |  |  |  |
| MG | 19 | 3,004 | less-affected<br>vs. more-affected | 29.8<br>(5.3) | 33.9<br>(5.6) | 4.1<br>(4.9) | 0.8 | 70.4 | 28.7 |

**Figure 5.**
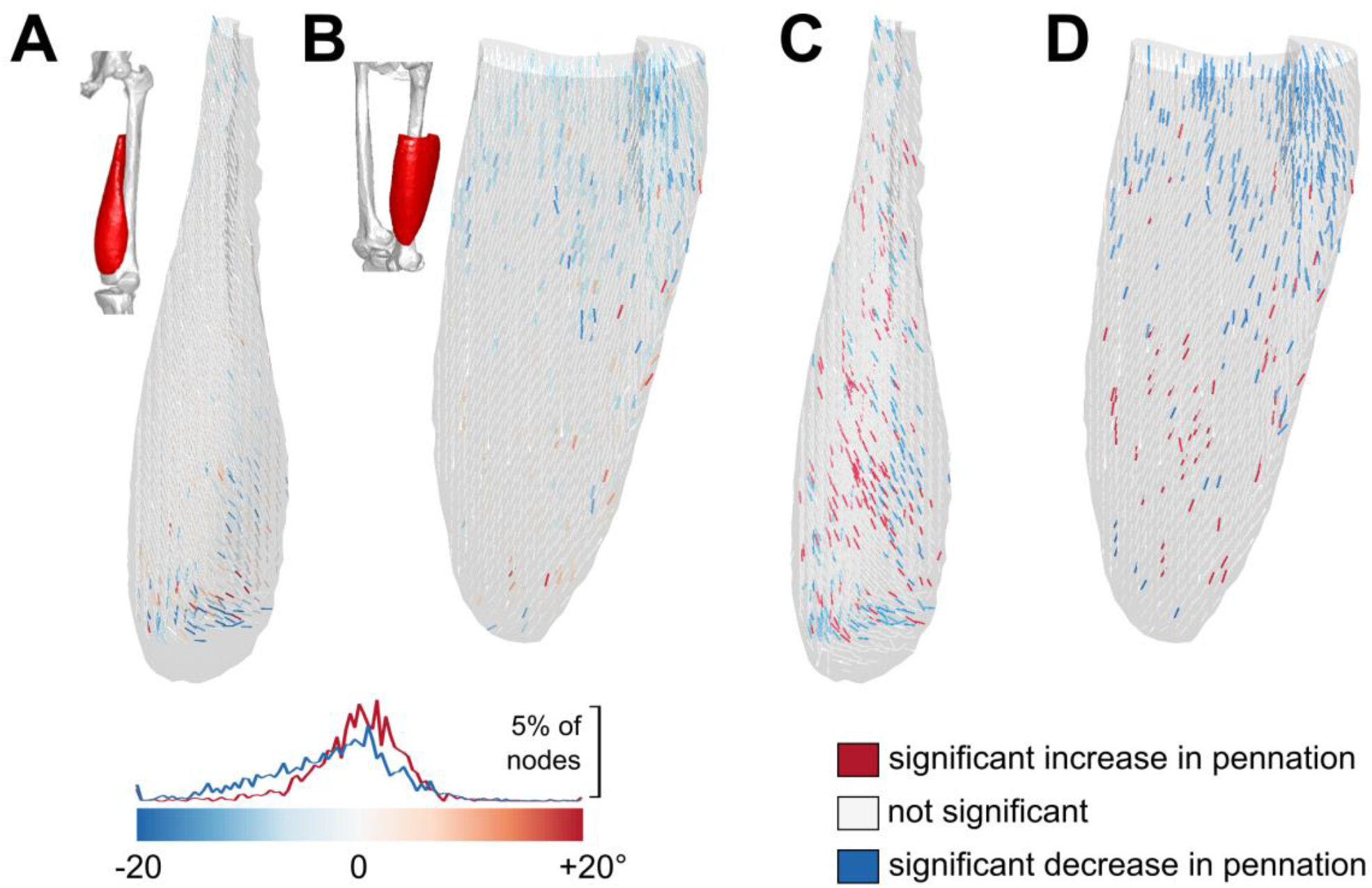
Effect of training on pennation angle of the vastus medialis (A,C) and vastus lateralis (B,D). In panels A and B, fibre orientation vectors are coloured according to their change in pennation with training (red/blue = larger/smaller pennation angle after training). The distribution of changes in pennation is plotted on top of the colour bar (red line for vastus medialis and blue line for vastus lateralis). In C and D, the fibre orientation vectors are coloured according to the statistical significance of the effect of training on pennation angle (red = significant increase, white = no significant change, blue = significant decrease).

### Effect of passive lengthening

Passive lengthening of the medial gastrocnemius resulted in a contraction of the surface in the middle of the muscle and an expansion near the proximal and distal ends, with larger changes in shape for larger changes in length (Fig. 6A and B). Both the contraction and the expansion were statistically significant for most of the superficial surface and both changes in length, demonstrating the ability of the methods to detect relatively small and heterogeneous changes in surface shape.

**Figure 6.**
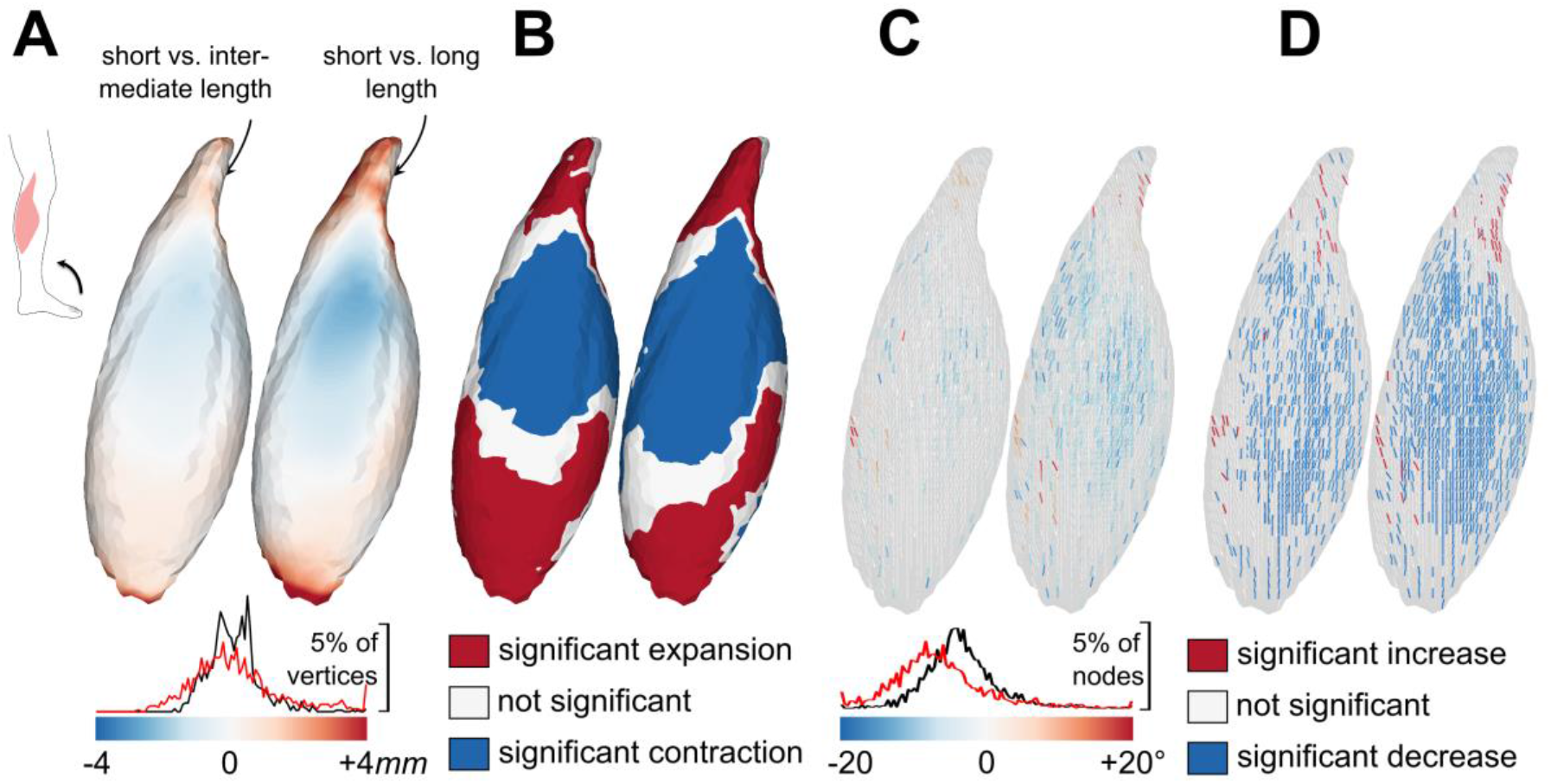
Effect of passive lengthening on muscle shape and architecture of the medial gastrocnemius muscle. A) The muscle surfaces are coloured according to the distance between the group-averaged surface at the shortest length to the surface at intermediate (left) and longest length (right; red/blue = local expansion/contraction). The distribution of distances is plotted on top of the colour bar (black line short to intermediate and red line short to long length). B) The muscle surfaces are coloured according to the statistical significance of changes in shape. C) Muscle architecture reconstruction with fibre orientation vectors coloured according to their change in pennation (red/blue = larger/smaller pennation compared to the shortest length). The distribution of changes in pennation is plotted on top of the colour bar (black line short to intermediate and red line short to long length). D) The fibre orientation vectors are coloured according to the statistical significance of the change in pennation. Each panel shows the same view on the superficial surface of the muscle. Changes in shape of the deep surface (not shown) were largely non-significant.

As expected, passive lengthening of the medial gastrocnemius oriented fibres more with the muscle’s long axis, resulting in an average decrease of pennation by 3.6° and 6.6° when the muscle was lengthened from a short to an intermediate and long length, respectively. The decrease in pennation was significant for 36% (short to intermediate length) and 53% (short to long length) of nodes.

### Effect of unilateral cerebral palsy

The smaller volume of the medial gastrocnemius on the more-affected side compared to the less-affected side of children with unilateral cerebral palsy (by 31% as reported previously) was visible as a substantial and statistically significant contraction across the whole superficial muscle surface (Fig. 7A and B). On average, the pennation angle was 4.1° larger on the more-affected side (Table 3), with the difference found to be statistically significant for 29% of nodes, mostly located in the central and proximal parts of the muscle (Fig. 7C and D).

**Figure 7.**
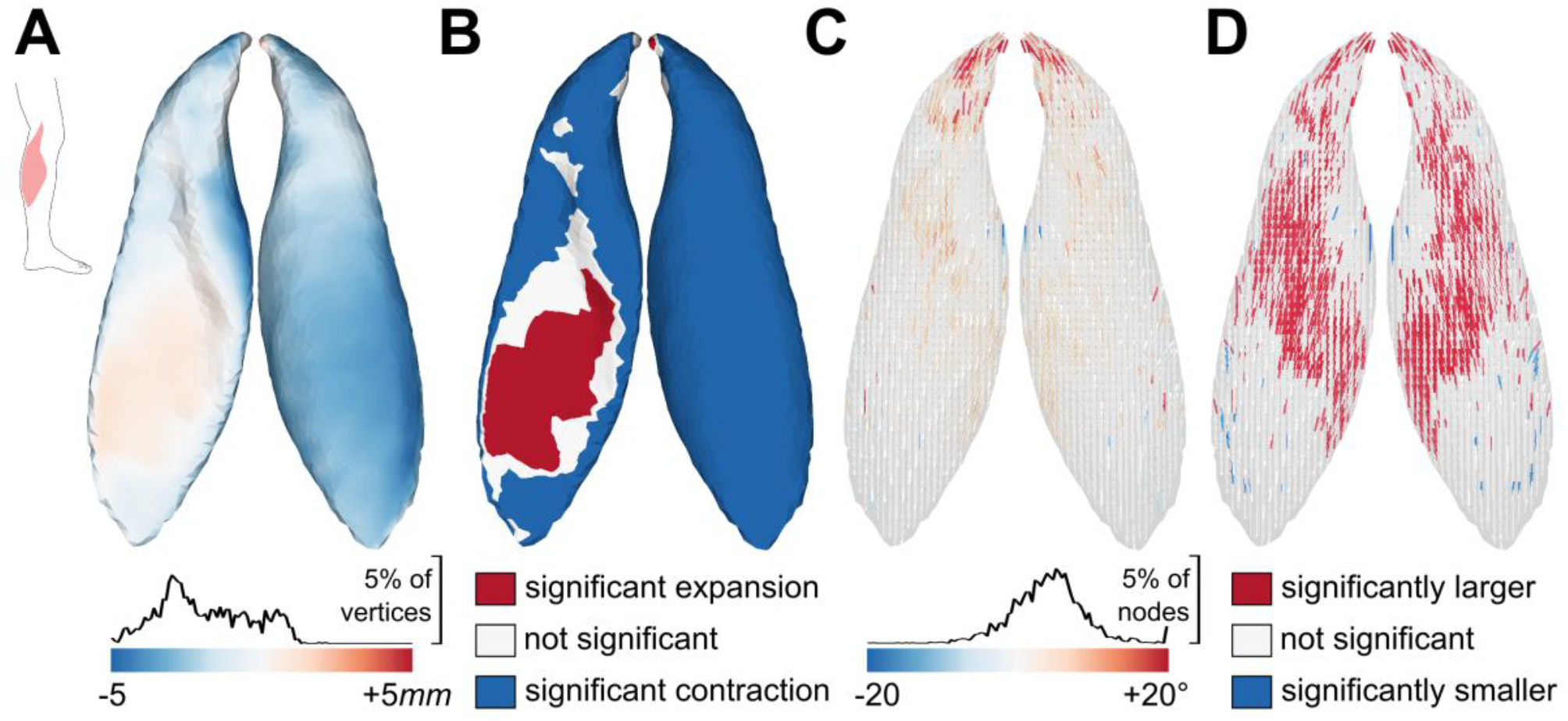
Comparison of medial gastrocnemius shape and architecture between the more- and less- affected side of children with unilateral cerebral palsy. A) Group-averaged muscle surface coloured according to the distance from the more-affected to the less-affected muscle (red/blue = local expansion/contraction). The distribution of distances is plotted on top of the colour bar. B) The muscle surfaces are coloured according to the statistical significance of differences in shape. C) Group-averaged fibre orientations coloured according to the difference in pennation angle between more- and less-affected muscles (red/blue = larger/smaller pennation in more-affected muscle). The distribution of changes in pennation is plotted on top of the colour bar D) The fibre orientation vectors are coloured according to the statistical significance of differences in pennation. Each panel shows two views of the same muscle.

## Discussion

This paper presents and makes available for general use a new framework for comprehensive analysis of changes in shape and architecture of human skeletal muscles from MRI and DTI data. Application of the methods to three datasets demonstrated the general use of the methods to quantify, visualise and statistically analyse complex changes in 3D shape and architecture. Strengths and weaknesses of the new methods will be discussed alongside the three key features of the proposed methodology: (1) point-to-point correspondence on surfaces and in volumes of muscles; (2) averaging of 3D muscle shapes and architectures; and (3) visualisation and statistical analysis of changes in shape and fibre orientations. Finally, areas of application and potential extensions of the new framework are discussed.

## Technical discussion

The ill-defined problem of point-to-point correspondence on complex 3D shapes is a general problem in, most notably, statistical shape modelling (Heimann and Meinzer, 2009). Various solutions have been proposed, of which the iterative closest point algorithms (Besl and McKay, 1992) or entropy-based particle systems (Cates et al., 2017) are probably the most popular. However, these algorithms only provide solutions for correspondence on surfaces, not inside volumes as was needed here to average fibre orientations inside muscles. Instead, we used non-rigid b-spline registration of distance maps of surfaces to define the smooth transformation of a volume that optimally aligns locations that are equidistant from the surface, which seems a reasonable solution for the correspondence problem on surfaces and in volumes. The b-spline transformation with a grid spacing of 20 mm was flexible enough to accurately transform the reference surface to all other surfaces (maximum residual error of 0.8 mm). By synthetically deforming muscles to simulate transversely uniform hypertrophy and passive lengthening, we showed very high accuracy of estimations of point-to-surface distances (root mean square errors <0.14 mm), and a good accuracy of point-to-point distances (0.3-1.3 mm; see Appendix 2 for details). While these tests demonstrate the adequacy of the methods to establish point correspondence on surfaces, correspondence of nodes inside the muscle is more difficult to evaluate quantitatively. The consequence of errors in correspondence is that fibre orientations are not only averaged across muscles (as intended), but also between locations. Large errors in correspondence would therefore lead to group-averaged models that do not preserve regional variation in muscle architecture. This did not appear to be a problem here: the group-averaged model of the vastus medialis clearly displayed the regional variation in architecture with twisting fibres along its length and oblique fibres near its distal insertion on the patella. This observation provides some evidence of the validity of the proposed solution for correspondence on surfaces and in volumes.

With the correspondence problem solved, vertex coordinates could simply be averaged across muscles to obtain group-averaged muscle surfaces. Averaging fibre orientations across muscles, was, however, more complex because diffusion tensors must be transformed into log-Euclidean domain before averaging (Arsigny et al., 2006). The architecture of the resulting group-averaged model was not only smoother than that of individual muscles, but also more complete: while diffusion tensors at 3-65% of nodes in individual muscles were deemed unrealistic, all group-averaged models were complete or nearly complete (Table 1). This is an important feature of group-averaged models, especially for applications in computational modelling for which sparse input data is problematic.

Group-averaged differences in shape were visualised by colour-coding surfaces by their signed point-to-surface distance between groups. Application of bootstrapping to determine statistical significance of local changes in shape revealed that strength training expanded the vastus medialis and vastus lateralis across the whole superficial surface with no clear evidence of regional variation in hypertrophy (Fig. 3). Changes at the deep surface were mostly non-significant, presumably indicating that the muscles were aligned mostly on the deep surface and that patterns of hypertrophy are best observed on the superficial surface. Shape analysis of muscles during passive lengthening demonstrated the ability to detect heterogeneous changes in shape with a small but significant contraction of the muscle surface centrally, while expanding in width and length. These effects were similar, but larger in magnitude, for larger changes in length (Fig. 6). When applied to the medial gastrocnemius of children with unilateral cerebral palsy (Fig. 7), it showed that muscles on the more-affected side are shorter, thinner and less wide than muscles on the less-affected side.

Bootstrapping was used to statistically analyse local differences in pennation angle between groups (Table 3). Despite some nodes reaching statistical significance in the strength training dataset, we interpreted the small mean change in pennation angle and the scattering of ‘significant’ changes throughout the muscle as evidence for little to no systematic effect of training on pennation angle. Application of this approach to passive lengthening muscles demonstrated that changes in pennation angle can be detected (Fig.6), but even with the largest change in pennation angle that was tested (on average 7°), statistical significance was reached for only 53% nodes. In the cerebral palsy dataset, changes in pennation were found to be statistically significant for 28% of nodes, clustered in the middle and proximal parts of the muscles. These findings suggest heterogeneous changes in pennation angle during passive lengthening and in muscles affected by cerebral palsy, but the sensitivity and robustness of the methods to determine statistically significant changes in fibre orientation requires further testing. Other statistical approaches like statistical parametric mapping may be used in future studies as well (Yamaguchi et al., 2021). Regardless of the statistical method, the relatively wide range of changes in pennation for all conditions tested (see distributions on colour bars in Fig. 5-7) suggest that it may prove difficult to reliably detect subtle changes in pennation angles between groups, unless large sample sizes or more accurate measurements of local fibre orientations are used.

## Future applications and extensions

The proposed framework enables investigations of skeletal muscle shape and architecture in more detail than possible with previous methods. The framework can naturally be applied to comprehensive investigations of muscle adaptation with, for example, childhood muscle growth, ageing or disease, or to evaluate the effect of other strength training protocols. The complete reconstructions of muscles could also be used to inform 3D continuum models of skeletal muscles to study the functional consequences of architectural adaptations. To date, the most detailed models use synthetically generated geometries and/or fibre architectures (Blemker and Delp, 2005; Choi and Blemker, 2013; Wakeling et al., 2020). Taking these conceptual models towards anatomically realistic analyses of subject-specific or population-averaged muscle function will require rich and accurate data on 3D muscle shape and architecture, which can be generated with the proposed framework. The methods might also be useful in the anatomical sciences, for example in comparative anatomy to compare muscle design between species, or to build anatomical atlases of muscle architecture.

The analyses conducted here focused on differences between group means, but the matrix of corresponding points can also be used to determine the major modes of variation in shape using principal component analysis, as is done in statistical shape modelling (Heimann and Meinzer, 2009). Statistical shape modelling and, by extension, statistical fibre orientation modelling (Peyrat et al., 2007) could further help identify and analyse group-level differences in 3D muscle shape and architecture; filter noisy measurements of muscle fibre orientations using information from a population; generate realistic muscle reconstructions from sparse imaging data; and identify potential relationships between muscle fibre orientations and muscle shape.

## Acknowledgements

I thank Prof Robert D Herbert for discussions about the methodology and for providing feedback on the manuscript, and Mr Junya Eguchi, Dr Arkiev D’Souza and A/Prof Jeanette Thom for help with data collection. The author acknowledges the facilities and scientific and technical assistance of NeuRA Imaging, a node of the National Imaging Facility, a National Collaborative Research Infrastructure Strategy (NCRIS) capability. No conflicts of interest, financial or otherwise, are declared by the author.

## Appendix 1: Imaging protocol for dataset 1 (strength training)

MRI scans of the left thigh were obtained immediately before the first training session and between 48 and 72 hours after the last training session on a 3T Philips Achieva TX MRI scanner (Philips Medical Systems, Best, The Netherlands). A 32-element cardiac coil, placed around the left thigh, was used for all acquisitions.

Participants lay supine with a wedge under the left knee to avoid compression of the posterior thigh against the MRI table from the weight of the leg. The calves were placed in U-shaped pieces of foam to maintain consistent positioning of the legs between days and participants. The angle between the left shank and the thigh was measured using a protractor and recorded to confirm consistent knee position between scans obtained before and after training.

The imaging protocol consisted of an mDixon and DTI scan of the left thigh with the following scan parameters:

- *mDixon*: 2-point dixon 3D-FFE sequence, FOV = 180 × 180 mm^2^, acquisition matrix = 180 × 180 (reconstructed to 192 × 192), TR/TE1/TE2 = 6.2/3.5/4.6 ms, flip angle = 6°, reconstructed voxel size = 0.94 × 0.94 × 1 mm^3^, number of slices = 320 and scan time of 355 seconds.
- *DTI*: EPI sequence, TR/TE = 8011/64, FOV = 180 × 180 mm^2^, flip angle = 90°, acquisition matrix = 92 × 90 (reconstructed to 112 × 112), reconstructed voxel size = 1.6 × 1.6 × 5 mm^3^, number of signal averages = 2, b-value = 500 s/mm^2^ (b0 with b = 0 s/mm^2^), number of slices = 50, number of gradient directions = 16 on a hemisphere, fat suppression: spectral attenuated inversion recovery (SPAIR) and scan time of 536 seconds. The DTI-scan was preceded by a B0-calibration scan.

## Appendix 2: Testing point correspondence

The procedures to determine point correspondence through registration of distance maps were tested by synthetically deforming muscle surfaces, and then comparing the known (true) distance between corresponding points to the estimated distances. Two metrics were tested:

- ***point-to-point distance:*** The Euclidean distance between corresponding vertices of the undeformed and deformed surface. The sign equals the sign of the projection of the displacement vector on the normal vector to the surface to approximate that negative/positive values indicate that the deformed vertex is inside/outside the undeformed surface, respectively.
- ***point-to-surface distance:*** the Euclidean distance between vertices on the deformed surface to *any* point on the undeformed surface. Negative/positive values indicate that the deformed vertex is inside/outside the undeformed surface, respectively. The Matlab function point2trimesh was used to calculate the point-to-surface distance ^1^.

The group-averaged vastus medialis and vastus lateralis before training were synthetically deformed to simulate hypertrophy. The muscles were first rotated to their local coordinate system using principal component analysis on the vertices of the muscles so that the first principal component indicates the long axis (x-axis) and the second (y-axis) and third principal components (z-axis) span the transverse plane. The centroid of all vertices was set as the origin of the local coordinate system. The y- and z-coordinates of all vertices were multiplied by 1.07 to simulate transversely uniform hypertrophy by 14.5%. Using similar procedures, the group-averaged medial gastrocnemius from the passive lengthening dataset was synthetically deformed with a pure shear deformation to simulate passive muscle lengthening. The point-to-point and point-to-surface distances were then estimated for all vertices through non-rigid registration of distance maps of the undeformed and deformed surface. The accuracy of the estimations was determined by calculating the root mean square error and Pearson’s correlation coefficient between the true and estimated distances.

For all three test cases, point-to-surface distances were estimated with very high accuracy (root mean square errors of 0.11-0.14 mm and correlation coefficients >0.96; Fig. S1). Point-to-point distances were estimated with lower accuracy than point-to-surface distances, but still with high accuracy (root mean square errors 0.28-1.23 mm and correlation coefficients >0.86). In conclusion, registration of distance maps gives an excellent estimation of point-to-surface distances and a good estimation of point-to-point correspondence, at least for the muscle deformations tested here.

**Figure S1.**
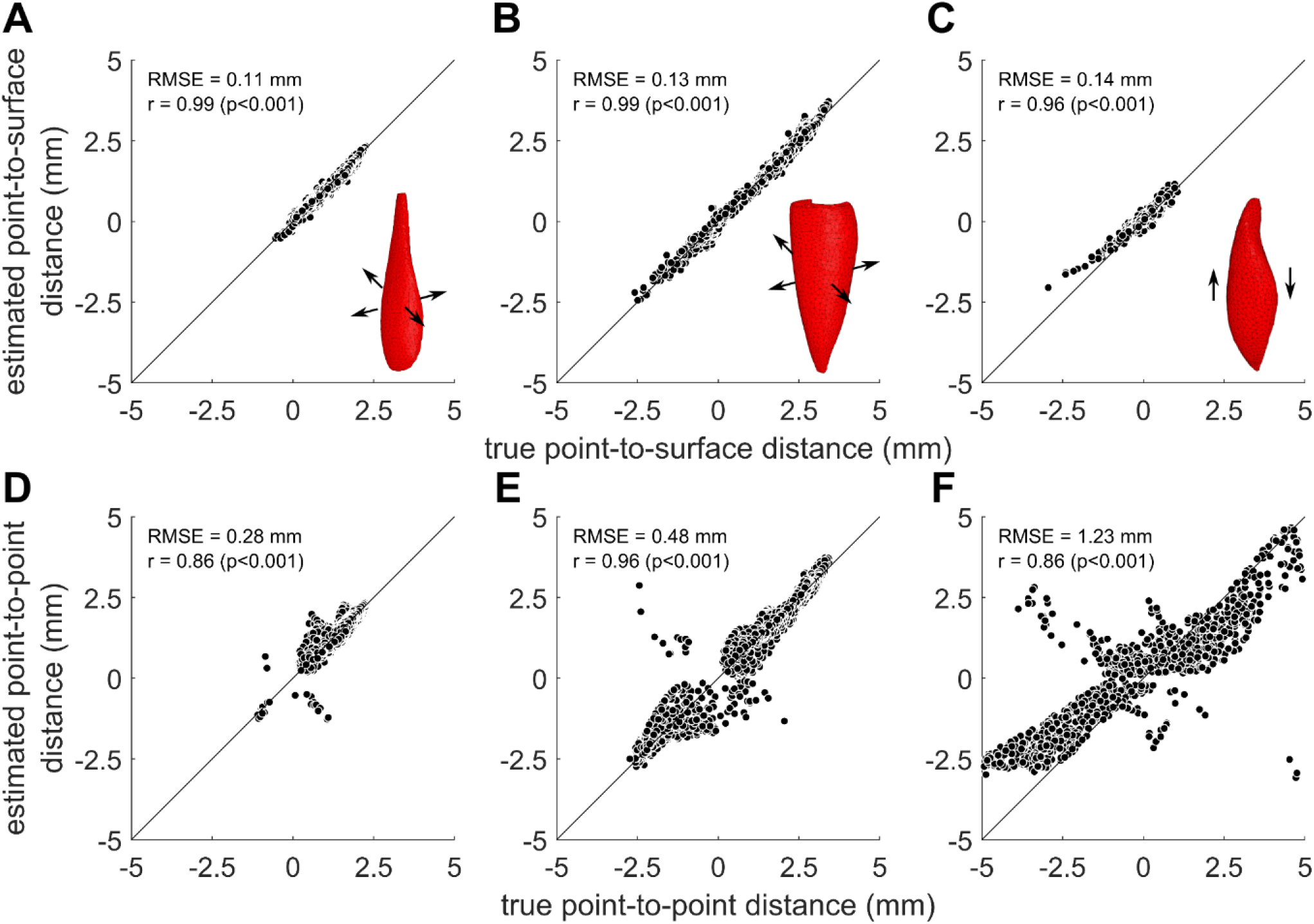
True vs. estimated point-to-surface (A-C) and point-to-point distances (D-F) between undeformed muscles and muscles that were synthetically deformed to simulate transversely uniform hypertrophy (A,D: vastus medialis, B,E: vastus lateralis) and passive lengthening (C,F: medial gastrocnemius). Insets show root mean square errors and Pearson’s correlation coefficient. The diagonal line is the line of identity.

## Footnotes

1 Daniel Frisch (2021). point2trimesh() — Distance Between Point and Triangulated Surface (https://www.mathworks.com/matlabcentral/fileexchange/52882-point2trimesh-distance-between-point-and-triangulated-surface), MATLAB Central File Exchange. Retrieved August 27, 2021.

